# Dual-purpose isocyanides produced by *Aspergillus fumigatus* contribute to cellular copper sufficiency and exhibit antimicrobial activity

**DOI:** 10.1101/2020.07.23.217810

**Authors:** Nicholas Raffa, Tae Hyung Won, Andrew Sukowaty, Kathleen Candor, Chengsen Cui, Saayak Halder, Mingji Dai, Julio Landero Figueroa, Frank C. Schroeder, Nancy Keller

**Affiliations:** Department of Medical Microbiology and Immunology, University of Wisconsin–Madison, Madison, Wisconsin, USA; Boyce Thompson Institute and Department of Chemistry and Chemical Biology, Cornell University, Ithaca, New York, USA; Department of Chemistry and Center for Cancer Research, Purdue University, West Lafayette, IN, USA; Agilent Metallomics Center of the Americas, Department of Chemistry, University of Cincinnati College of Arts and Science, Cincinnati, OH 45221, USA; Department of Pharmacology and System Biology, University of Cincinnati College of Medicine, Cincinnati, OH 45267, USA; Department of Bacteriology, University of Wisconsin—Madison, Madison, Wisconsin, USA

## Abstract

The maintenance of sufficient but non-toxic pools of metal micronutrients is accomplished through diverse homeostasis mechanisms in fungi. Siderophores play a well-established role for iron homeostasis; however, no copper-binding analogs have been found in fungi. Here we demonstrate that in *Aspergillus fumigatus* isocyanides derived from the *xan* biosynthetic gene cluster (BGC) bind copper, impact cellular copper content, and have significant metal-dependent antimicrobial properties. *xan* BGC-derived isocyanides are secreted and bind copper as visualized by a chrome azurol S (CAS) assay and inductively coupled plasma-mass spectrometry (ICP-MS) analysis of *A. fumigatus* intracellular copper pools demonstrated a role for *xan* cluster metabolites in the accumulation of copper. *A. fumigatus* coculture with *A. nidulans, Candida albicans* and a variety of pathogenic bacteria establish copper-dependent antimicrobial properties of *xan* BGC metabolites including inhibition of laccase activity. Similarly, inhibition of *Pseudomonas aeruginosa* by low concentrations of the *xan* isocyanide xanthocillin was copper-dependent. Other metals also reduced xanthocillin’s antimicrobial properties, but less efficiently than copper. As variations of the *xan* BGC exist in other filamentous fungi, we suggest that xanthocillin-like natural products represent a first example for fungal small molecules that serve to maintain copper sufficiency and mediate interactions with competing microbes.

**Significance:** Metal homeostasis is an integral part of metabolism for any organism. A vast array of small molecules are already known to mediate metal homeostasis in fungi and bacteria; however, unlike their bacterial counterparts, to-date there are no known fungal small molecules that function to maintain copper homeostasis. Discovery of copper binding small molecules produced by *A. fumigatus* gives insight into mechanisms other than the extensively studied copper transporters or metalloproteins for how fungi can regulate copper. This has important ecological implications as securing scarce nutrients is central for fitness and survival. Additionally, studying this mechanism in *A. fumigatus* provides a basis for investigation of copper regulation pathways in other fungi.

## Introduction

*Aspergillus fumigatus* is a ubiquitous, filamentous fungus that exists in the environment as a saprophyte, feeding on dead organic matter. It is also the primary causative agent of invasive aspergillosis, a disease with a mortality rate of over 90% in the most severe cases (1, 2). In both environments *A. fumigatus* is a common microbiome constituent (e.g. respiratory tract biofilms with pathogenic bacteria including *Pseudomonas aeruginosa* (3), staphylococci (4) and *Stenotrophomonas* spp. (5) or soil communities composed of numerous fungi, bacteria and protists (6, 7)). *A. fumigatus* secondary metabolites are low-weight, bioactive chemicals that can mediate fungal success in these diverse environments (8).

Secondary metabolites help *A. fumigatus* overcome several hurdles, one of which is regulation of trace element homeostasis, assuring sufficiency for use as cofactors in enzymes while simultaneously limiting their toxic effects (9, 10). For example, intracellular and extracellular siderophores are produced by *A. fumigatus* as important components of acquisition and storage of iron (11, 12). Similar to iron, copper is a micronutrient required for function of a variety of enzymes including superoxide dismutases (SOD)(13), cytochrome c oxidase (14), and proteins involved in reductive iron uptake (FetC)(12). While essential, copper also participates in Fenton chemistry by reacting with hydrogen peroxide to generate toxic hydroxyl radicals (15) and can also displace other metal cofactors (e.g. Fe, Zn) in enzymes rendering them inert (16). Thus, intracellular copper content must be precisely balanced.

Small molecule regulators of copper homeostasis have been identified in bacteria but have yet to be described in fungi. Methanobactins are shown to be essential for copper uptake in methanotrophic bacteria, necessary for the functioning of copper-dependent methane monooxygenase (17, 18). The small molecule yersiniabactin is a virulence factor produced by several pathogenic strains of *E. coli* and acts to mitigate the toxicity of copper by binding to it, but also ensures adequate supply of copper to cells (19– 21). One class of small molecules commonly involved in copper binding are isocyanides (also known as isonitriles)(22). For instance, the bacterium *Streptomyces thioluteus* produces and secretes SF2768, an isocyanide that has been demonstrated to be responsible for copper uptake (23) and the entomopathogenic bacterium, *Xenorhabdus nematophila*, produces the isocyanide virulence factor rhabduscin which has been shown to inhibit the insect immune defense copper-dependent laccase enzyme (24).

The first fungal isocyanide <u>b</u>iosynthetic <u>g</u>ene <u>c</u>luster (BGC) was recently identified in *A. fumigatus* (25). Using chemical and genetic tools, here we demonstrate that *xan* BGC metabolite(s) are secreted and bind copper, affecting both cellular copper levels in *A. fumigatus* and interactions with other microbes. Overexpression of the *xan* BGC inhibits endogenous and exogenous laccase activity and demonstrates metal-dependent broad-spectrum antimicrobial activity. Xanthocillin, one of the isocyanide products of the *xan* BGC, recapitulates the *xan* BGC copper-dependent antibacterial activity at micromolar concentrations. This is the first characterization of copper-binding small molecules produced by fungi which, further, supports a role for these natural products in copper homeostasis and microbial competition.

## Results

### Overexpression of *xanC* results in a copper-dependent pigmentation defect in *Aspergillus fumigatus*

Prior evidence suggests that the *xan* BGC is regulated by the copper-binding transcription factors AceA and MacA (25), therefore we first wanted to determine the response of the *xanC* mutants to different levels of copper. To achieve this, we inoculated plates of GMM supplemented with various amounts of copper with the *xan* cluster overexpression mutant (Figure 1A). The OE::*xanC* mutant displayed a growth defect across all levels of copper and had a spore pigmentation phenotype when grown on media containing 0 μM and 5 μM CuSO_4_ that is remediated as the copper concentration increases. Since this phenotype is identical to the pigmentation defect of the wild type grown under copper limiting conditions this indicates that overexpression of the *xan* BGC is causing a copper deficiency. We speculated the poor growth could be due to decreased functionality of copper-dependent proteins (e.g. SODs and laccases) including the laccase-mediated production of the green spore pigment DHN melanin (Figure 1B). Indeed a laccase activity assay showed reduced activity in the OE::*xanC* strain relative to wild type at all levels of copper tested (Figure 1C). This likely explains the spore pigmentation defect of this strain at 5 μM CuSO_4_ as two copper-dependent laccases are required for production of DHN-melanin in *A. fumigatus* (26). Taken together, these data suggest that the metabolites produced by the *xan* BGC are inhibiting copper-dependent processes in *A. fumigatus*.

**Figure 1.**
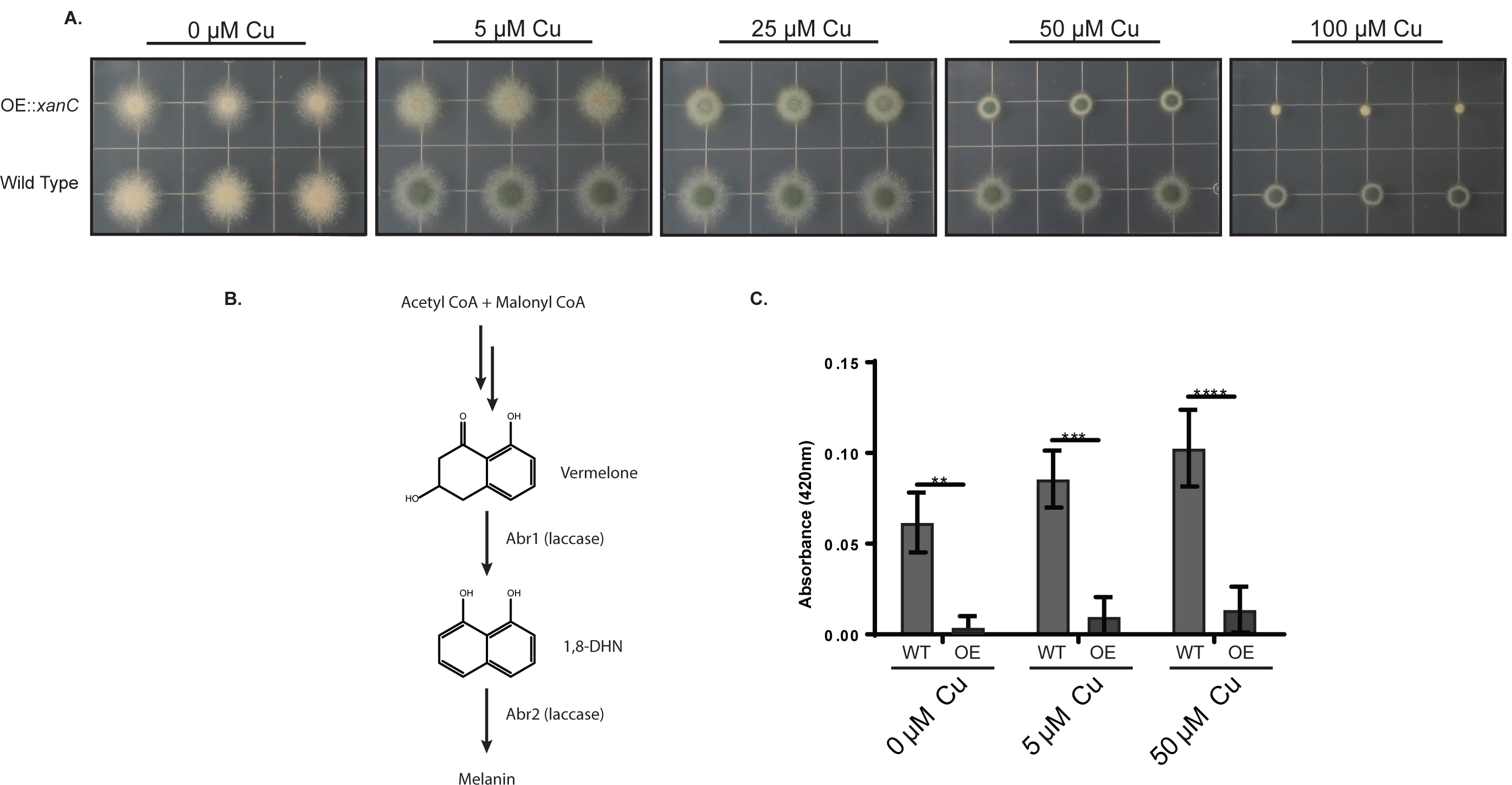
The response of *xan* cluster mutants to copper stress yields a copper-dependent pigmentation phenotype. A.) Growth of OE::*xanC* mutants on solid GMM supplemented with CuSO_4_. B.) Biosynthesis of DHN-melanin by *A. fumigatus* C.) ABTS laccase activity assay of *xanC* mutants when grown on solid media supplemented with CuSO_4_. Statistical analyses were performed using Student’s t-test and shown is the mean with SEM of three replicates. OE = OE::*xanC*, WT = wild type. *P<0.05, **P<0.01, ***P<0.001, ****P<0.0001.

### Gene deletions in the *xan* BGC confirm their role in the biosynthetic pathway

Our earlier study led to a proposed *xan* BGC biosynthetic pathway (25). To validate this proposed pathway as well as to gain knowledge of the types of metabolites synthesized by the *xan* BGC, we created a series of double mutants by deleting each *xan* gene in the OE::*xanC* background (Figure 2A). LC-MS analysis of the double mutants confirmed the role of XanB as the isocyanide synthase as there were no detectable isocyanides in the OE::*xanC*Δ*xanB* mutant (Figure 2B-H and Table S3). XanG was confirmed to dimerize the isocyanide monomer, since when *xanG* was deleted we observed increased accumulation of a monomeric isocyanide species and abolishment of the production of isocyanide dimers. Analysis of its tandem mass spectrometry (MS/MS) spectra and molecular formulae suggested that the monomer represented a dehydrogenated formyl tyrosine (**3**, Figure 2H, and Table S4). This result suggested that tyrosine is converted into the monomer in a two-step sequence analogous to bacterial PvcA-PvcB isocyanide synthase (27), followed by dimerization of the monomer by XanG. Deletion of *xanE* confirmed the role of XanE in the methylation of the isocyanides, resulting in decreased total isocyanide production and methylated xanthocillin derivatives. Deletion of *xanA* yielded an identical phenotype to the OE::*xanC* strain suggesting that it is not involved in the conversion of the isocyanide moiety to its *N*-formyl derivative, even though XanC shares homology with previously studied isocyanide hydratases (Figure S2). XanD is a small DUF4149 protein, and deleting it results in similar levels of the isocyanide derivatives to the OE::*xanC* strain, indicating that *xanD* is not involved in biosynthesis of any of the known *xan* BGC metabolites. Deletion of the hypothetical protein, XanF, results in decreased isocyanide production relative to wild type, suggesting that XanF may improve efficiency of isocyanide production. BLAST analysis of XanF indicates the presence of a methyltransferase domain, suggesting that it could potentially work with XanE to methylate the xanthocillin. However, it is also possible that in the process of deleting XanF the flanking genes, XanG and/or XanE, were disrupted during transformation. Overall, chemical analysis shows differential accumulation of *xan* BGC products which we then used as a tool to investigate which metabolites were responsible for the growth inhibition and copper phenotype.

**Figure 2.**
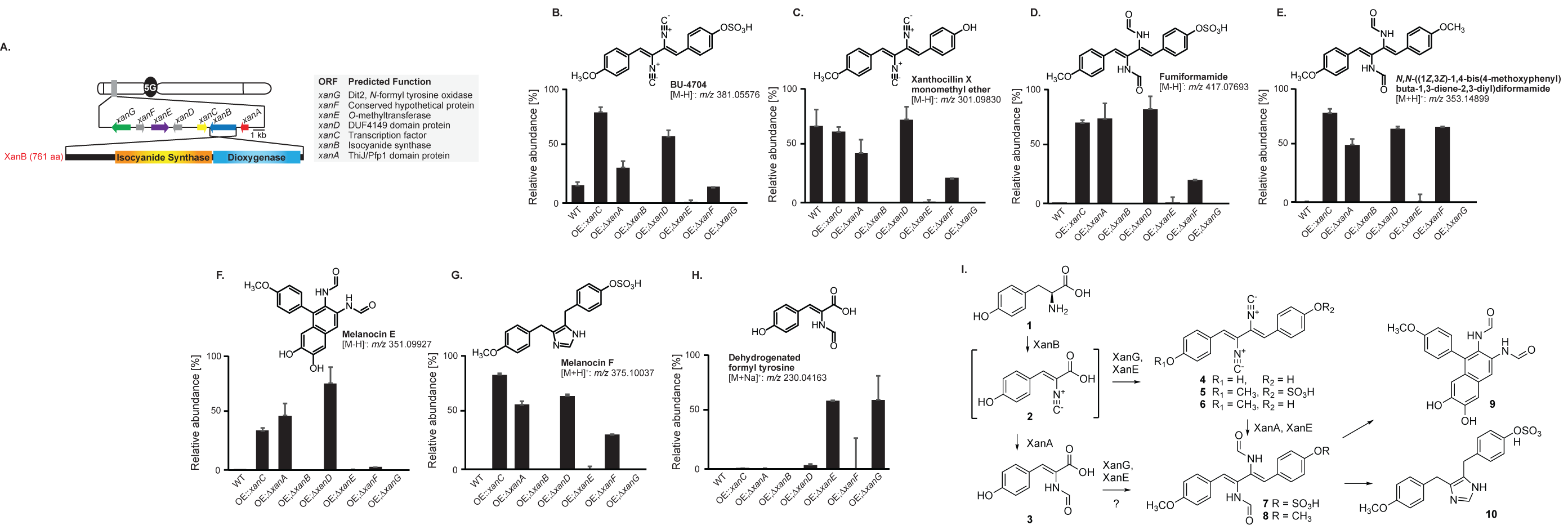
*xan* gene cluster and putative functions of encoded proteins. A.) The *xan* gene cluster responsible for the production of xanthocillin derivatives contains seven genes. *xanA* (ThiJ/Pfp1 domain protein with homology to isocyanide hydratases, red); *xanB* (two domain isocyanide synthase-dioxygenase, blue); *xanC* (C6 transcription factor, yellow); *xanD* (DUF4149 domain protein, grey); *xanE* (O-methyltransferase, purple); *xanF* (Conserved hypothetical protein, grey); *xanG* (P450 monooxygenase homology to the yeast enzyme Dit2, green). B-H). Relative abundances, as determined by HPLC-HRMS in ESI^+^ or ESI^-^-mode, of xanthocillin derivatives in WT, OE::*xanC*, and a series of double mutants in which individual *xan* genes were deleted in the OE::xanC background. WT-wild type, OE;Δ*xanX* - OE::*xanC*;Δ*xanX*. I.) Putative biosynthesis of xanthocillin derivatives in *A. fumigatus*. Results from the current study further support the *xan* BGC biosynthetic pathway proposed in our previous study (25). Isocyanide containing intermediate (**2**) produced from tyrosine (**1**) by XanB is dimerized to form xanthocillin (**4**) and related isocyanides (BU-4704, **5**, and xanthocillin X monomethyl ether, **6**) by XanG. A dehydrogenated formyl tyrosine (**3**) likely represents a shunt metabolite derived from **2**. Xanthocillins are converted into the corresponding formyl derivatives (fumiformamide, **7**, and *N,N*-((1Z,3Z)-1,4-bis(4-methoxyphenyl)buta-1,3-diene-2,3-diyl)diformamide, **8**) by XanA and other isocyanide hydratases, which can undergo additional transformations to produce melanocin E (**9**) and melanocin F (**10**). Methylation of hydroxyl moieties in xanthocillin derivatives was likely introduced by XanE.

### Isocyanide accumulation is associated with extracellular copper-binding and loss of spore pigmentation

Chemical profiling of the *xan* BGC mutants allowed for grouping into those that produced high levels of isocyanides (OE::*xanC*, OE::*xanC*;Δ*xanA*, and OE::*xanC*;Δ*xanD*) and those that produced less or no detectable amount (OE::*xanC*;Δ*xanB*, OE::*xanC*;Δ*xanE*, OE::*xanC*;Δ*xanF*, and OE::*xanC*;Δ*xanG*). Considering that some bacterial isocyanides are secreted and bind copper (18), we hypothesized that *xan* BGC isocyanides would function similarly. To assess secretion, we performed a CAS assay using the *xan* BGC mutants. The OE::*xanC*, OE::*xanC*;Δ*xanA* and OE::*xanC*;Δ*xanD* – the three mutants that produced highest levels of isocyanides – showed similar levels of extracellular copper-chelating ability (Figure 3A, FigureS2), whereas the other double mutants resembled more closely the copper-binding phenotype of the wild type strain. This suggested a role for *xan*-derived isocyanides as secreted copper chelators.

**Figure 3.**
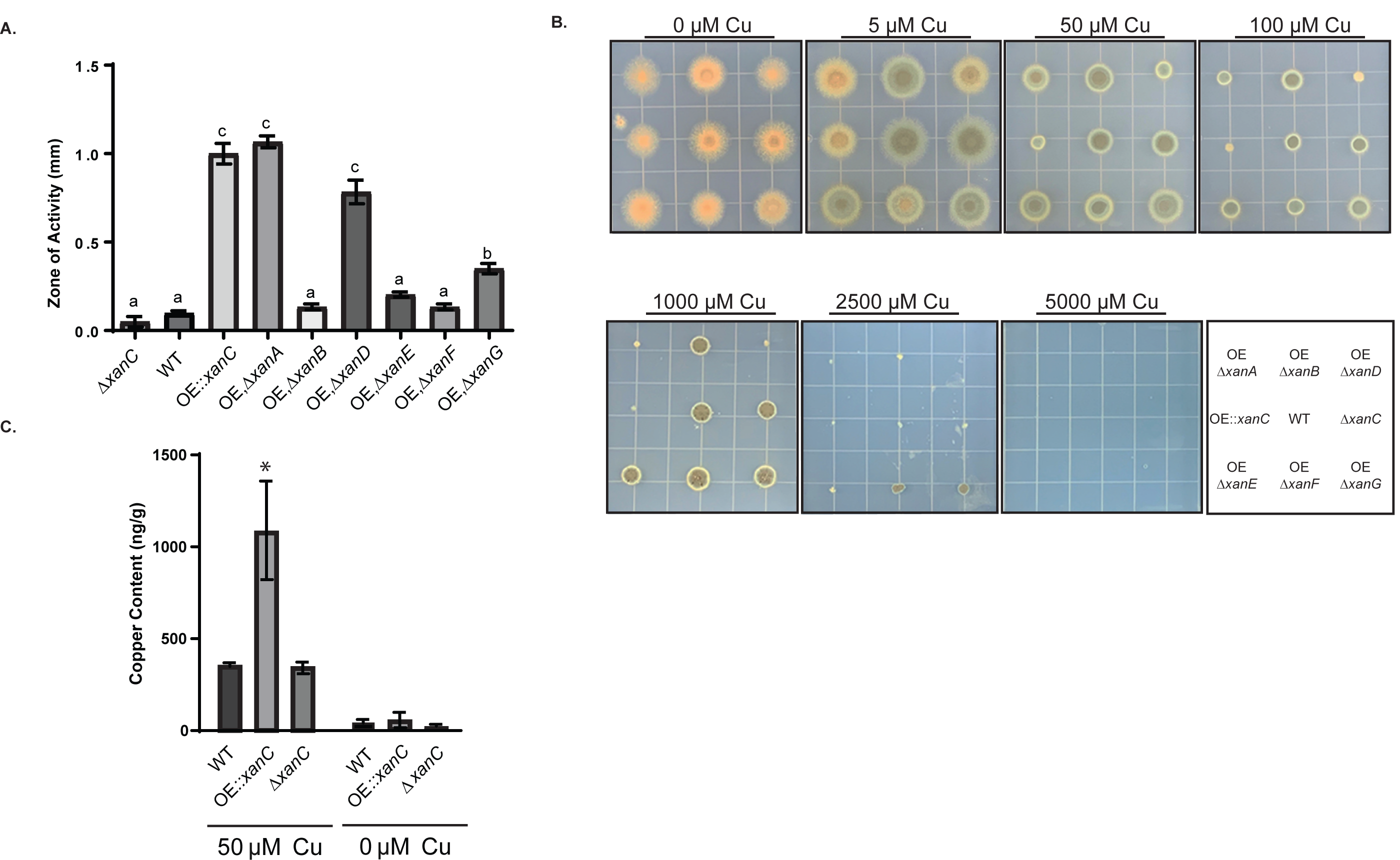
Involvement of *xan* cluster metabolites in copper secretion and uptake. A.) Quantification of the zone of activity of CAS assay of *xan* BGC double mutants. Measurements and error bars are representative of the mean of three replicates and standard error, respectively. Letters indicate statistically similar groups via an ANOVA. B.) Growth of the *xan* BGC double mutants on GMM supplemented with CuSO_4_. Bottom right-Strain location key. WT = wild type, OE, Δ*xanA =* OE::*xanC*;Δ*xanA*, OE, Δ*xanB =* OE::*xanC*;Δ*xanB*, OE, Δ*xanD =* OE::*xanC*;Δ*xanD*, OE, Δ*xanE = OE::xanC;*Δ*xanE*, OE, Δ*xanF =* OE::*xanC;*Δ*xanF*, OE, Δ*xanG =* OE::*xanC*;Δ*xanG*. C.) Cellular copper content of *xan* cluster mutants grown in liquid shake culture, either lacking copper or supplemented with 50 μM CuSO_4_, as determined by ICP-MS. Error bars represent SEM of three replicates and statistical analysis was performed using an ANOVA. ‘*’ indicates P < 0.05 relative to wild type.

To further investigate whether isocyanide production was linked to copper-related phenotypes, we tested the responses to copper of each of the double mutants. The OE::*xanC*;Δ*xanA* and OE::*xanC*;Δ*xanD* mutants displayed a growth and pigmentation defect compared to the OE::*xanC* strain (Figure 3B). Interestingly, the OE::*xanC*;Δ*xanF* mutant produced both isocyanides and their derivatives, yet did not exhibit the same copper binding phenotype either in the CAS assay or with regards fungal growth. Comparison of the metabolomes of these four strains shows that OE::*xanC*;Δ*xanA*, OE::*xanC*;Δ*xanD*, and OE::*xanC* all produce higher total amounts of isocyanides relative to OE::*xanC*;Δ*xanF* (Figure 2). These data suggest that the *xan* BGC-derived isocyanide are primarily responsible for the pigmentation and extracellular copper-binding phenotype.

### *xan* BGC metabolites are associated with the accumulation of cellular copper

In the bacterium *Streptomyces thioluteus*, the chalkophore SF2768 is secreted, binds copper and is then taken up by the cells (23). Based on the finding that *xan* BGC metabolites are secreted and bind copper, we asked if *xan* BGC metabolites could directly affect copper uptake. We grew the wild type, OE::*xanC*, and Δ*xanC* strains in liquid culture either supplemented with 50 □M copper or without copper supplementation and analyzed their intracellular copper concentrations by ICP-MS. We found that intracellular copper concentrations of the OE::*xanC* mutant had elevated levels of intracellular copper relative to wild type in the 50 □M copper treatment (Figure 3C). There was no detectable difference in copper accumulation in the no copper treatment. Thus, it appears that *xan* BGC metabolites are associated with an increase in cellular copper accumulation.

### *xan* BGC mutants inhibit pigmentation in *Aspergillus nidulans* and exhibit broad-spectrum antimicrobial properties when grown in coculture

Recent studies have shown that fungal secondary metabolites are synthesized and/or protect the producing fungus during encounters with bacteria and fungi (28), such as induction of the *A. fumigatus* fumicycline A BGC during interactions with the soil bacterium *Streptomyces rapamycinicus* (29). Further, considering such secondary metabolites have been examined as potential antimicrobials, we wanted to determine if the *xan* BGC metabolites demonstrate antimicrobial properties or otherwise influence microbial interactions.

In view of the inhibitory effect of *xan* BGC metabolites on laccase activity (Figure 1C), we cocultured wild type *A. fumigatus* and the OE::*xanC* mutant with *A. nidulans* to determine if the *xan* metabolites inhibit *A. nidulans* laccase activity. The *A. nidulans* laccase Ya is required for the green pigment of its spores, and when Ya is inactive or deleted, the spores are yellow (30), yielding an easily scored phenotype. Figure 4A shows that *A. nidulans* spores remained yellow along the border with the OE::*xanC* mutant compared to the wild type control, where *A. nidulans* presented normal spore pigmentation. This pigmentation defect was rescued when the coculture was performed on media supplemented with excess CuSO_4_.

**Figure 4.**
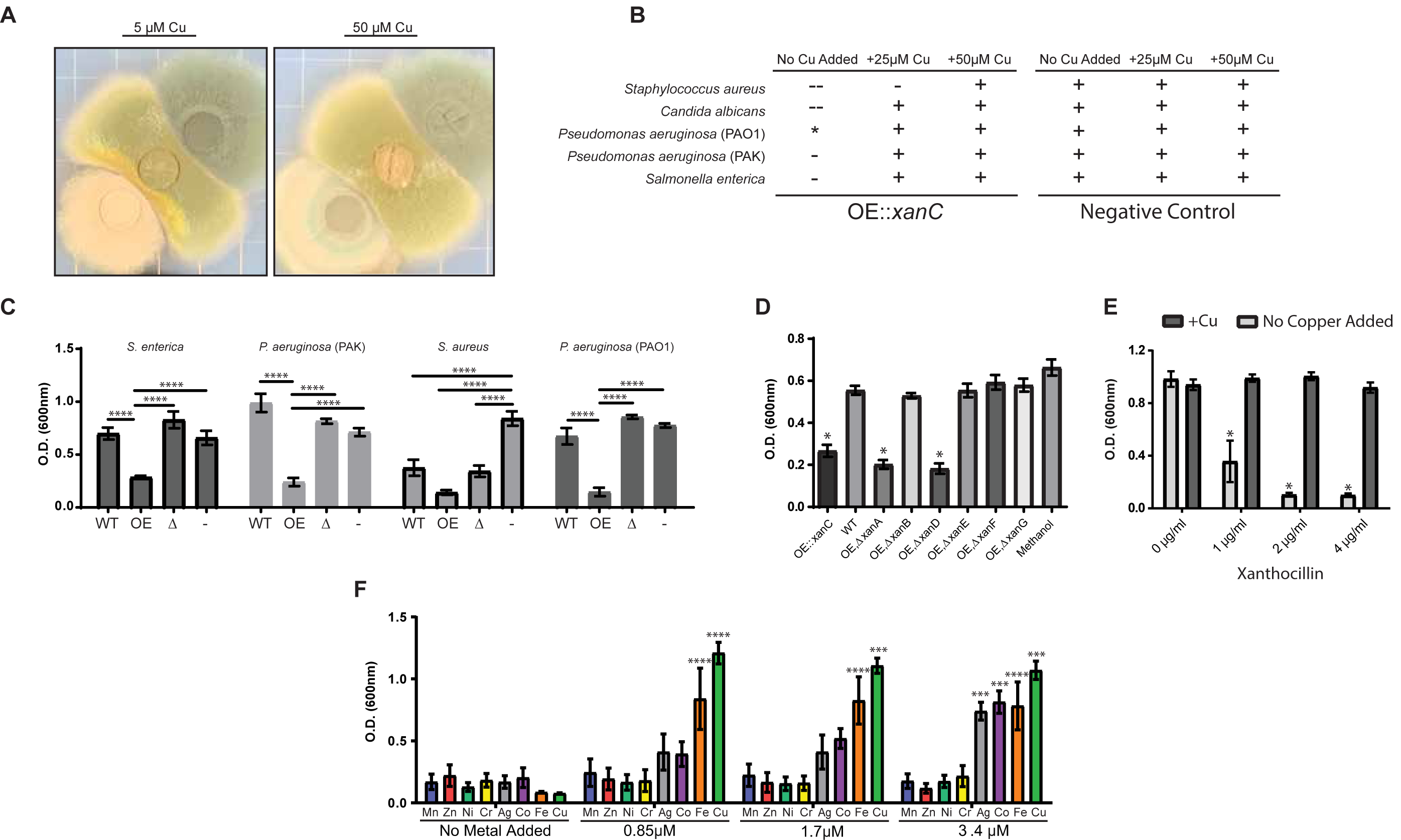
*xan* cluster metabolite affect growth and copper-dependent processes of microorganisms in a metal-dependent fashion. A.) Coculture of *Aspergillus nidulans* (middle of plate) with wild type *A. fumigatus* (top right of plate) and the OE::*xanC* mutant (bottom left of plate) on solid GMM supplemented with CuSO_4._ B.) Cocultures of *xan* mutants with microorganisms grown on solid LB. Growth inhibition of the indicate microbe was qualitatively assessed by visually inspecting the relative sizes of the zone of inhibition around the agar plugs. ‘**--**’ = inhibition, ‘**-**’ = partial inhibition, ‘**+**’ = no inhibition, ‘**\***’ = inhibits pigments production C.) Growth assay of microorganisms challenged with WT, OE::*xanC*, and Δ*xanC* extracts. OE = OE::*xanC*, WT = wild type, Δ = Δ*xanC*, ‘**-**’ = methanol (vehicle control). D.) Growth assay of *P. aeruginosa* (PAO1) challenged with *xan* BGC double mutant extracts. WT = wild type, OE, Δ*xanA =* OE::*xanC*;Δ*xanA*, OE, Δ*xanB =* OE::*xanC*;Δ*xanB*, OE, Δ*xanD =* OE::*xanC*;Δ*xanD*, OE, Δ*xanE = OE::xanC;*Δ*xanE*, OE, Δ*xanF =* OE::*xanC;*Δ*xanF*, OE, Δ*xanG =* OE::*xanC*;Δ*xanG*. E) *P. aeruginosa* PAO1 challenged with purified synthetic xanthocillin with and without 3.5 μM CuSO_4_ supplemented. F.) *P. aeruginosa* PAO1 challenged with 1μg/ml purified synthetic xanthocillin and supplemented with heavy metals. Statistical analyses were performed using an ANOVA, comparing to WT (D), comparing within each xanthocillin treatment (E), or comparing to growth at 0 μM of the indicated metal treatment (F). Error bars represent SEM of experiments performed in triplicate. *P<0.05, **P<0.01, ***P<0.001, ****P<0.0001.

To determine if the *xan* BGC metabolites have antimicrobial properties, actively growing cultures of *A. fumigatus* mutants were cocultured with representative microbes. We observed significant inhibition of *Staphylococcus aureus, Candida albicans*, and *Salmonella enterica*, as well as loss of blue pigmentation in *Pseudomonas aeruginosa* (PAO1) next to the OE::*xanC* mutant (Figure 4B, Figure S3). The pigmentation and growth phenotypes were abolished when the media was supplemented with copper for all of the microbes tested, suggesting that *xan* BGC metabolites have copper-dependent antimicrobial properties, for example by inhibiting copper uptake or copper-dependent processes in other microbes.

We next tested the antimicrobial activity of *A. fumigatus* extracts against *Staphylococcus aureus, Salmonella enterica, Pseudomonas aeruginosa* (PAO1), and *Pseudomonas aeruginosa* (PAK). There was a consistent inhibition of growth of all of the microbes challenged with the OE::*xanC* extract (Figure 4C). As with confrontations with actively growing fungus (Figure 4B, Figure S3), we found that addition of copper was able to abolish the growth defect of *P. aeruginosa* when challenged with the OE::*xanC* extract (Figure S4). To narrow down the *xan* BGC metabolites responsible for this activity, we performed the assay again using extracts from the double mutants. There was consistent antibacterial activity in extracts from the double mutants that produce the highest concentration of isocyanides, OE::*xanC*;Δ*xanD* and OE::*xanC*;Δ*xanA*, similar to that of the OE::*xanC* (Figure 5D). These data suggest that it is indeed the isocyanides that are responsible for the inhibition of pigment production and antimicrobial properties.

### Pure synthetic xanthocillin has potent copper-dependent antibacterial activity

The patterns of *A. fumigatus* mutant growth (Figure 1A), the copper-binding properties of mutant secretions (Figure 3A) and antimicrobial activities (Figures 4) all support a view that the *xan* BGC-derived isocyanides are the sole bioactive molecules from this pathway. To confirm this hypothesis, we compared inhibitory activities of pure xanthocillin prepared by total synthesis (31) and several *N*-formyl derivatives of the isocyanides (which no longer feature an isocyanide moiety) against *Pseudomonas* spp. We found that only xanthocillin inhibited bacterial growth (Figure 4E and Figure S5) at a MIC of 3.5 □M (1 □g/ml), placing it at par with current MICs of gentamicin (2 □g/ml), ceftazidime (1 □g/ml), and imipenem (1.5 □g/ml)(32). As with the crude *xan* BGC extracts, the inhibitory effect of xanthocillin was lost with increasing levels of copper. Since we previously showed that copper is able to abolish the inhibitory properties of the *xan* mutant extracts and when grown in coculture, we wanted to determine if the antimicrobial activity of xanthocillin can be abolished with the addition of copper as well. We repeated the MIC assay and found that addition of a 1:4 molar ratio of xanthocillin to copper was able to completely abolish the antimicrobial activity of the isocyanide, rescuing the growth to wild type levels (Figure 4E). This indicates that the antimicrobial activity of the xanthocillin is dependent on the absence of high copper concentrations.

### Pan-metal remediation of isocyanides antimicrobial activity

Metal chelating small molecules often bind more than one metal, for instance yersiniabactin binds to copper, nickel, and iron (19, 33), and pyoverdine, the siderophore produced by *Pseudomonas*, has been shown to bind copper and nickel as well. To determine whether metals other than copper can abolish *xan* metabolite antimicrobial activity, we added different heavy metals to cultures of *Pseudomonas aeruginosa* PAO1 challenged with OE::*xanC* extract. Addition of nickel, cobalt, and iron were able to rescue the growth defect to the same levels of the copper control, whereas manganese, molybdenum, and zinc were not effective (Figure S4). We next asked if these same metals could abolish the antibacterial activity of pure xanthocillin and found that addition of iron, cobalt, and silver were also able to mitigate the inhibitory properties of the extract in the case of *P. aeruginosa* PAO1 (Figure 4F). These data suggest that the *xan* BGC isocyanides bind specific metal ions, including but not limited to copper.

## Discussion

Copper homeostasis is a critical requirement for microbial success in all environments. Whereas tight transcriptional regulation of copper importers, exporters, and storage proteins has been well characterized in many fungi (34), including *A. fumigatus* (35), until now no small molecule(s) has been found to contribute to copper biology in any fungal system. Our findings are the first to identify secondary metabolites that impact cellular copper content in fungi. This work potentially adds a eukaryotic member to the small but growing number of bacterial systems that synthesize copper binding small molecules as a mechanism for copper uptake (12, 18, 33). We find *xan* BGC isocyanides provide *A. fumigatus* with a competitive edge in coculture challenge and suggest that they may have a role as antimicrobials in the environment and could potentially be repurposed for clinical uses.

Through the creation of *xan* BGC double mutants, we provide compelling evidence that the isocyanide moieties of this cluster bind copper and increase cellular copper content. Only the three mutants that accumulated the highest concentration of isocyanides (OE::*xanC*;Δ*xanA*, OE::*xanC*;Δ*xanD* and OE::*xanC*) showed positive CAS assay results and laccase pigmentation defects (Figures 1 and 3). Although we did not find an increase of copper in the OE::*xanC* strains under copper starvation conditions (Figure 3C), we speculate that our analytical methods may not have been sensitive enough to record a small but significant difference in copper levels under these limiting conditions. Together, our data supports the notion that *xan* BGC metabolites are secreted, bind copper, inhibit activity of copper-requiring enzymes, and possibly contribute to copper sufficiency.

Use of *xan* BGC mutants and purified *xan* BGC metabolites strongly suggest that the isocyanide moieties of the *xan* BGC metabolites were responsible for their antimicrobial activities. The isocyanide producing mutants that showed high CAS activity, *A. fumigatus* pigmentation, and growth defects were the same strains to inhibit or alter bacterial growth. Isocyanides also inhibited pigmentation in *P. aeruginosa* (PAO1), which we hypothesize is due to an interference with the copper-pyocyanin complex (36). Furthermore, only xanthocillin but not *N*-formyl isocyanide derivatives significantly inhibited *Pseudomonas* growth (Figure 4E-F, S5). Antimicrobial or laccase inhibiting properties were abolished by increasing copper concentrations in media (Figure 4). Additional heavy metals such as silver, cobalt, and iron were able to rescue the growth defect of the bacteria challenged with xanthocillin, suggesting that the isocyanides bind to a subset of metal cations, with xanthocillin itself having the highest affinity for copper under the tested conditions. Overall, this indicates that isocyanides could be used by *A. fumigatus* to compete with other microbes by either depriving competitors of essential, scarce nutrients or by directly inhibiting metal-dependent processes. This supposition is strongly supported by a previous study where coculture of the bacterium *Streptomyces peucetius* with *A. fumigatus* induced synthesis of several *xan* cluster metabolites associated with bacterial inhibition (at the time the *xan* cluster had not been identified, (37)).

Furthermore, we have confirmed the roles of most Xan proteins in the biosynthesis of the *xan* BGC metabolites, solidifying the proposed biosynthetic pathway presented in a previous study (25). Two *xan* proteins that did not affect biosynthesis under the tested conditions were XanD and XanA. XanD is a protein of unknown function; however, XanA shares homology with previously studied isocyanide hydratases (38). In bacteria, isocyanide (isonitrile) hydratases convert isocyanides to their less toxic *N*-substituted formamides (39). Hence we speculated that the *OE::xanC*J*xanA* strain would be sicker than *OE::xanC* by virtue of producing more isocyanides and less *N*-formyl derivatives. This was not the case. One possible explanation for the apparent lack of involvement of XanA in this pathway is that there are six other putative isocyanide hydratases in the *A. fumigatus* genome. This redundancy suggests that at least one of these other proteins is able to convert the *xan* isocyanides to their *N*-formyl derivative, thus compensating for the absence of XanA.

In conclusion, we here report a small molecule(s) produced by a fungus that preferentially binds copper. This work adds a eukaryotic system to the many reported bacterial species known to use small molecules to bind copper, possibly functioning in copper uptake, and in related interactions with other organisms. Although this work focused on *A. fumigatus*, it seems likely that similar molecules play analogous roles in other fungi, as the *xan* BGC is conserved in several fungal species (25). As with *A. fumigatus*, many of these fungi possess more than one isocyanide synthase-containing BGC. The characterization of these additional BGCs should provide exciting new insights in the functions of isocyanides in the microbial world.

## Materials and Methods

All strains and primers used in this study are listed in the supplemental material (Table S1, S2). Details of construction of mutants, fungal growth conditions, physiological assays, and chemical analysis are provided in supplemental data. Statistical analyses were performed using an ANOVA or Student’s t-test and the program GraphPad Prism.

## Acknowledgements

Partial funding for this work was provided by the National Institutes of Health under grant R01GM112739 to FCS and NPK, R35GM128570 to MD, and T32GM008349. The funders had no role in study design, data collection and interpretation, or the decision to submit the work for publication.

## Supplemental Text

### Materials and Methods

#### Mutant construction

Double-joint PCR was used to generate DNA constructs for transformation described previously (1). DNA constructs containing the *pyrG* or *argB* selectable marker fused to 1kB homologous regions flanking the gene of interest that insert via homologous recombination. Flanks were amplified with 20bp overlaps using primers designed using SeqBuilder (DNASTAR, Madison, WI). Selectable markers were amplified from either the pJW24 (*pyrG*) (2) or pJMP4 (*argB*) (3) plasmids. Deletion mutants were constructed by whole-gene deletion and overexpression mutants were created by inserting a constitutively active *A. nidulans gpdA* promoter upstream of the ATG translation start site of a gene. The DNA constructs were transformed into the *pyrG* auxotroph TFYL 80.1 via PEG transformation described previously (1). For double mutants, the double auxotroph TFYL 44.1 (*pyrG*^*-*^ and *argB*^*-*^) was first transformed with either the overexpression or deletion *xanC* construct and upon successful transformation, further transformed with a DNA deletion construct containing the *argB* marker. Resulting transformants were then screened via PCR using the 5’ forward flank and the marker reverse primer. Positive mutants were further screened via southern blot analysis using P-32 labelled 1kB flanking regions described previously to confirm single integrations (Figure S1).

#### Fungal Growth Conditions and Physiological Assays

Unless otherwise specified, *Aspergillus* strains were grown on glucose minimal medium (GMM)(4). *Aspergillus* strains were activated by streaking out on a plate of GMM from a glycerol stock. Spore stocks were generated by harvesting spores using 0.01% Tween 20 in ddH_2_O and gentle scrubbing using a L-shaped spreader. Spores were then filtered through sterile MiraCloth (Calbiochem) to remove debris and washed twice with Millipore-filtered water. Spore stocks were counted using a hemocytometer and diluting to the desired concentration. For testing response to varying levels of copper, GMM was prepared as described previously (5) without copper or EDTA. After autoclaving, copper or BCS was supplemented. Plates were inoculated with 2 μl of spore suspension and the plates grown at 37 °C for 72 hours. For liquid shake cultures, 50 ml of GMM was inoculated with 1×10^6^ spores/ml and incubated at 37 °C, shaking at 250 rpm for 48 hours. Lyophilized mycelia were obtained by filtering cultures through MiraCloth, freezing in liquid nitrogen, and lyophilizing overnight.

#### Chrome Azurol S Assay

Chrome azurol S (CAS) assay plates were made by adding 5 mL of sterile phosphate buffer (26 g/L KH_2_PO_4_ and 62 g/L Na_2_HPO_4_ – 7H_2_O) and 100 ml of 10x CAS (0.5 mM CuSO_4_, 0.5025 mM Chrome Azurol S (Sigma-Aldrich), 1.05 mM HDTMA) to 1 L of molten GMM. Upon solidifying, a sterile razor was used to remove half of the CAS media. 10 ml of warm molten GMM was then inoculated with 1×10^7^ spores and poured into the vacant space. The plates were then allowed to solidify and incubated for 5 days at 37 °C in the dark. The zone of activity was quantified by measuring the distance between the growing mycelia and the edge of the Cu-CAS complex with a ruler.

#### Laccase Activity Assay

Approximately 1⨯10^7^ spores were inoculated on small plates of GMM for 48 hours at 37°C in the dark. 2,2’-azino-bis(3-ethylbenzothiazoline)-6-sulphonic acid (ABTS) (Sigma-Aldrich) stock solution was prepared by dissolving ABTS in 0.1 M sodium acetate buffer (pH 4.5). Plates were flushed with 10 ml of 1 mM ABTS and incubated at ambient temperature for 24 hours. 200 μl aliquots of supernatant were transferred to a 96-well plate and measured the absorbance at 420 nm. Data were quantified and normalized to the negative control, uninoculated plates with the appropriate level of copper, flushed, and incubated with 1 mM ABTS.

#### Metabolite Extraction

Strains were grown as an overlay culture, inoculating 10 ml of warm, molten GMM (containing 8 g/L agar) with 1×10^6^ spores and pouring onto a plate containing 20 ml of solidified GMM. Upon solidification, the plates were then incubated for 5 days at 37 °C. The plates were frozen using liquid nitrogen and lyophilized. The lyophilized samples were extracted with 20 ml of ethyl acetate-methanol (9:1) for 1.5 h with vigorous stirring. Extracts were filtered over cotton, evaporated to dryness, and stored in 4-ml vials. Crude extracts were suspended in 0.5 ml of methanol and centrifuged to remove insoluble materials, and the supernatant was analyzed by UHPLC-HRMS.

#### HPLC-HRMS Analytical Methods and Equipment Overview

High-resolution HPLC-MS (HPLC-HRMS) was performed on a ThermoScientific-Dionex Ultimate 3000 UHPLC system equipped with a diode array detector and connected to a ThermoScientific Q Exactive Orbitrap mass spectrometer operated in electrospray positive (ESI^+^) or electrospray negative (ESI^−^) ionization mode. An Agilent Zorbax RRHD Eclipse XDB-C18 column (2.1 × 100 mm, 1.8 μm particle diameter) was used with acetonitrile (organic phase) and 0.1% formic acid in water (aqueous phase) as solvents at a flow rate of 0.5 mL/min. A solvent gradient scheme was used, starting at 2% organic for 1 min, followed by a linear increase to 100% organic over 14 min, holding at 100% organic for 2.5 min, decreasing back to 2% organic for 0.1 min and holding at 2% organic for the final 1.4 min, for a total of 18 min.

#### ICP-MS

The spores and mycelia were analyzed by ICP-MS after acid digestion.100 µl of concentrated trace metal grade nitric acid, 50 µl of 18 MΩ water and 25 ul of 500 ng/ml of scandium used as internal standard solution were added to the samples in a metal free 5 ml conical vial. After the digestion was carried out in a heating block at 90 °C for 2 hours with venting every 20 minutes. The samples were brought to 2.5 ml with 18 MΩ water. The samples were then analyzed for total Cu content in an Agilent 7500 ICP-MS system with a Cetac ASX-520 auto sampler contained in an acrylic box. The ICP-MS system was configured with a Micromist nebulizer, a double pass Scott spray chamber held at 2 °C, a 2.5 mm torch with platinum shield torch, and nickel sample and skimmer cones. The instrument was run with 3.5 ml/min of helium in energy discrimination mode. The external calibration method was used with a calibration range of 0.05 to 25 ng/ml. The mass of the samples (200-800 µg) needed for the quantification of Cu was calculated by measuring the phosphorous content in the digested samples according to (6).

#### Total Synthesis of Xanthocillin

Synthetic xanthocillin was prepared in 10 steps from commercially available 4-hydroxybenzaldehyde by following the approach developed by Tatsuta and co-workers (7). The NMR and HRMS data of the synthetic sample (Supplemental Figure S6) match with the reported ones. Pure synthetic xanthocillin sample was obtained after column chromatorgraphy for the corresponding biological evaluations.

#### Xanthocillin synthesis

To a solution of xanthocillin silly ether (prepared using the reported approach;^7^ 12 mg, 0.023 mmol) in THF (1 mL) at 0 °C under an atmosphere of argon was added TBAF/AcOH (90 uL, 1: 1/mol: mol, 0.09 mmol, premixed) dropwise. The reaction mixture was stirred at 0 °C for 30 min and then warm up to room temperature for 10 min. The reaction mixture was purified as it was by column chromatography (silica gel, 10-35% EtOAc in hexane) to provide xanthocillin (5 mg, 76% yield) as a yellow solid. *R*_*f*_= 0.27 (33% EtOAc in hexane); IR (film) 3275, 2131, 1593, 1515, 1434, cm^−1^; ^1^H NMR (500 MHz, CD_3_OD) δ 7.73 (d, *J*=8.5, 4H), 7.00 (s, 2H), 6.88 (d, *J*=8.5, 4H); ^13^C NMR (125 MHz, CD_3_OD) δ 174.6, 161.0, 133.0, 128.7, 125.0, 116.9, 116.7; HRMS (ESI): calcd for C_18_H_13_N_2_O_2_ (M+H)^+^ 289.0977, found 289.0972. The ^1^H and ^13^C NMR spectra of synthetic xanthocillin were recorded on a 500 MHz NMR spectrometer in CD_3_OD, with residual CH_3_OH signal(s) as the internal reference. Chemical shifts are reported as δ values (ppm). The solvent peaks are set as follows: CH_3_OH at d 3.31 and d 49.0 ppm for ^1^H and ^13^C NMR. Chemical shifts are reported as follows: s = singlet, d = doublet. IR spectra were taken on an FT-IR spectrophotometer. High-resolution mass spectra (HRMS) were measured by the ESI method.

Silica gel was used for flash column chromatography with mixed ethyl acetate (EtOAc) and hexane as the eluting solvents. Yields refer to chromatographically and spectroscopically (^1^H NMR) homogeneous materials. Anhydrous THF was freshly distilled from sodium benzophenone ketyl under argon. Other reagents were obtained commercially and used as received.

#### Cocultures and Growth Conditions

*Candida albicans, Staphylococcus aureus, Salmonella enterica, Pseudomonas aeruginosa* (PAK), and *Pseudomonas aeruginosa* (PAO1) were activated from glycerol stock onto plates of lysogeny <u>b</u>roth (LB) or <u>y</u>east <u>p</u>eptone <u>d</u>extrose (YPD), streaking for single colonies. A single colony was then selected and inoculated into 5 ml of liquid LB or YPD, shaking at 37 °C overnight. Coculture experiments were performed by transferring GMM agar plugs of an actively growing overlay fungal culture to solid LB containing 16 g/L agar. An overlay of the bacteria/fungi was generated by suspending microorganisms to a final O.D._600_ of 0.05 in 10 ml of molten top agar and dispensing onto the LB containing the agar plugs. The plates were then incubated overnight at either 37 °C in the case of the bacterial strains or 30 °C in the case of the *Candida albicans*. Zones of inhibition were visually evaluated to determine inhibition of growth. For cocultures involving *Aspergillus nidulans*, GMM containing various levels of copper was inoculated with 1×10^7^ spores of *A. nidulans* and preincubated for one day at 37 °C. The plates were then inoculated with 1×10^7^ spores of the *xan* mutant strains and incubated for an additional three days at 37 °C. For 96-well plate assays, overnight cultures were diluted to an O.D._600_ of 0.05 in liquid LB and 190 μl was dispensed into a well containing 10 μl of the corresponding extract/xanthocillin. The plate was then incubated at 37 °C. Growth was tracked by checking the O.D._600_ at 48 hours. For assays involving the addition of metals, filter-sterilized 1000x stocks were diluted in LB prior to dispensing into 96-well plates.

**Supplemental Figure S1.**
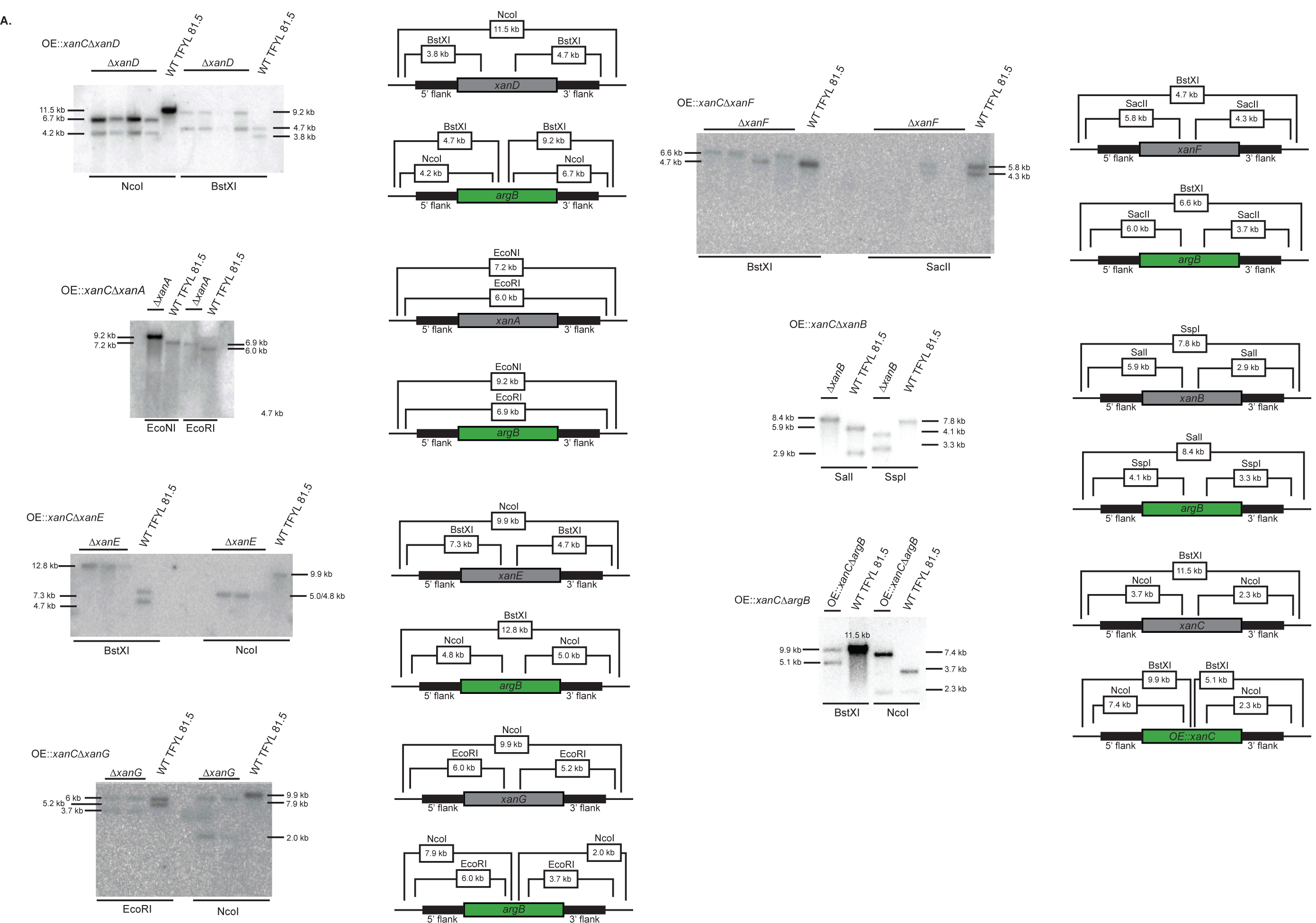
Southern Blot Confirmation analysis of mutants used in this study. Shown is the blot with labelled digest band sizes (Left) and a schematic of the digest including restriction enzymes used with the expected band sizes (Right).

**Supplemental Figure S2.**
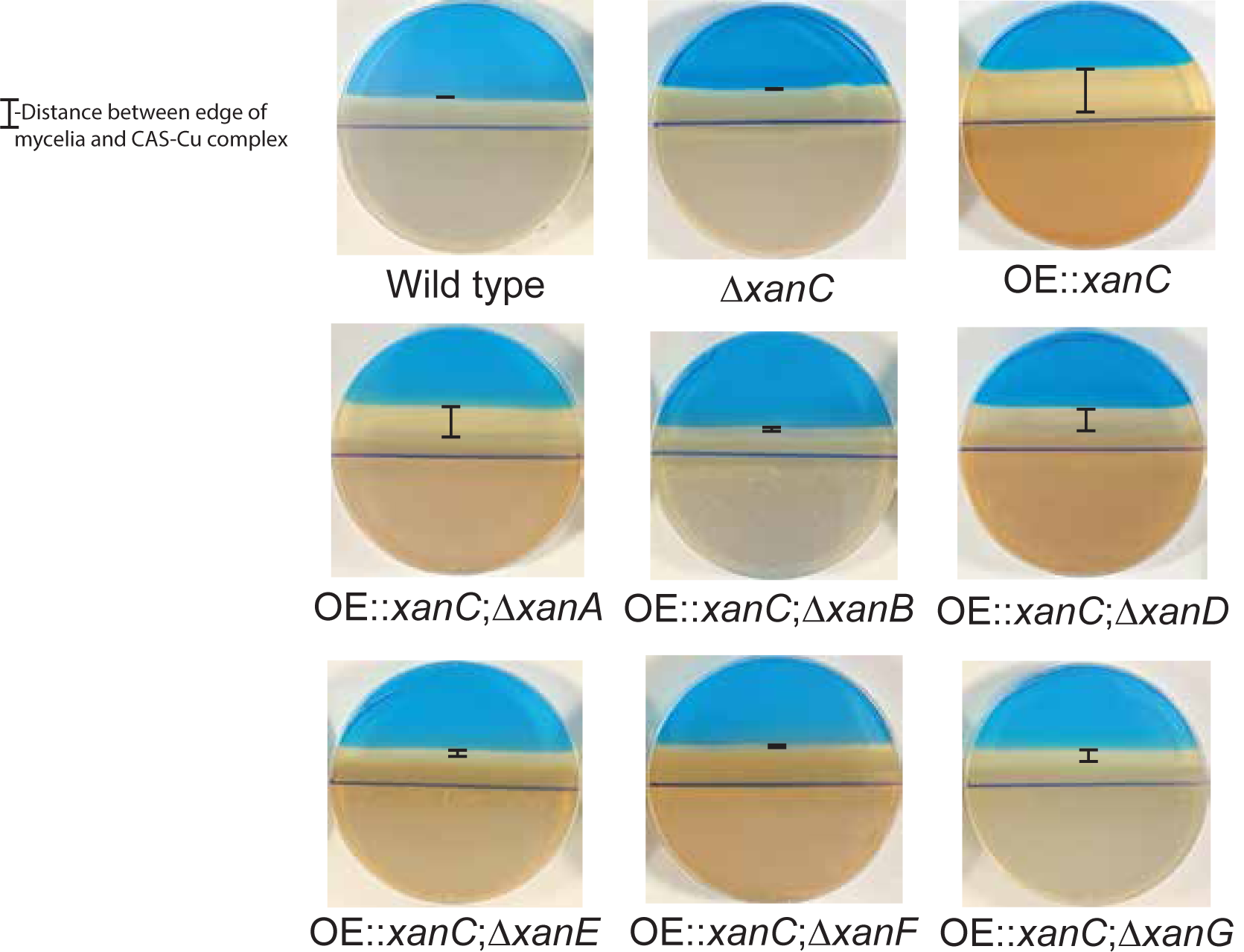
CAS assay for detection of secreted copper-binding molecules. Representative images of the CAS assay results. Measured distances of the zone of activity are indicated. Zone of activity is defined as the distance between the edge of the growing mycelia and the CAS-Cu complex.

**Supplemental Table S1.** Strains used in this study.

| Strain | Brief Genotype | Genotype | Reference |
| --- | --- | --- | --- |
| TFYL 81.5 | WT | pyrG1, argB1, fumiargB, fumipyrG, ΔakuA::mluc | Throckmorton 2016 |
| TFYL 84.1 | ΔargB | pyrG1, argB1, fumipyrG, ΔakuA::mluc | Lim 2018 |
| TFYL 80.1 | ΔpyrG | pyrG1, argB1, fumiargB, ΔakuA::mluc | Lim 2018 |
| TFYL 45.1 | ΔpyrGΔargB | pyrG1, argB1, ΔakuA::mluc | Throckmorton 2016 |
| TNLR 11.3 | OE::xanCΔargB | pyrG1, argB1, parapyrG::gpdA(p)::AFUA_5G02655; ΔakuA::mluc | This Study |
| TNLR 9.1 | ΔxanC | pyrG1, argB1, fumiargB, ΔAFUA_5G02655::parapyrG; ΔakuA::mluc | Lim 2018 |
| TNLR 1.2 | OE::xanC | pyrG1, argB1, fumiargB, parapyrG::gpdA(p)::AFUA_5G02655; ΔakuA::mluc | Lim 2018 |
| TNLR 12.1 | OE::xanCΔxanA | pyrG1, argB1, parapyrG::gpdA(p)::AFUA_5G02655; ΔAFUA_5G02670::fumiargB; ΔakuA::mluc | This Study |
| TNLR 13.2 | OE::xanCΔxanB | pyrG1, argB1, parapyrG::gpdA(p)::AFUA_5G02655; ΔAFUA_5G02660::fumiargB; ΔakuA::mluc | This Study |
| TNLR 20.5 | OE::xanCΔxanD | pyrG1, argB1, parapyrG::gpdA(p)::AFUA_5G02655; ΔAFUA_5G02650::fumiargB; ΔakuA::mluc | This Study |
| TNLR 17.5 | OE::xanCΔxanE | pyrG1, argB1, parapyrG::gpdA(p)::AFUA_5G02655; ΔAFUA_5G02640::fumiargB; ΔakuA::mluc | This Study |
| TNLR 18.6 | OE::xanCΔxanF | pyrG1, argB1, parapyrG::gpdA(p)::AFUA_5G02655; ΔAFUA_5G02630::fumiargB; ΔakuA::mluc | This Study |
| TNLR 19.2 | OE::xanCΔxanG | pyrG1, argB1, parapyrG::gpdA(p)::AFUA_5G02655; ΔAFUA_5G02620::fumiargB; ΔakuA::mluc | This Study |

**Supplemental Table S2.**
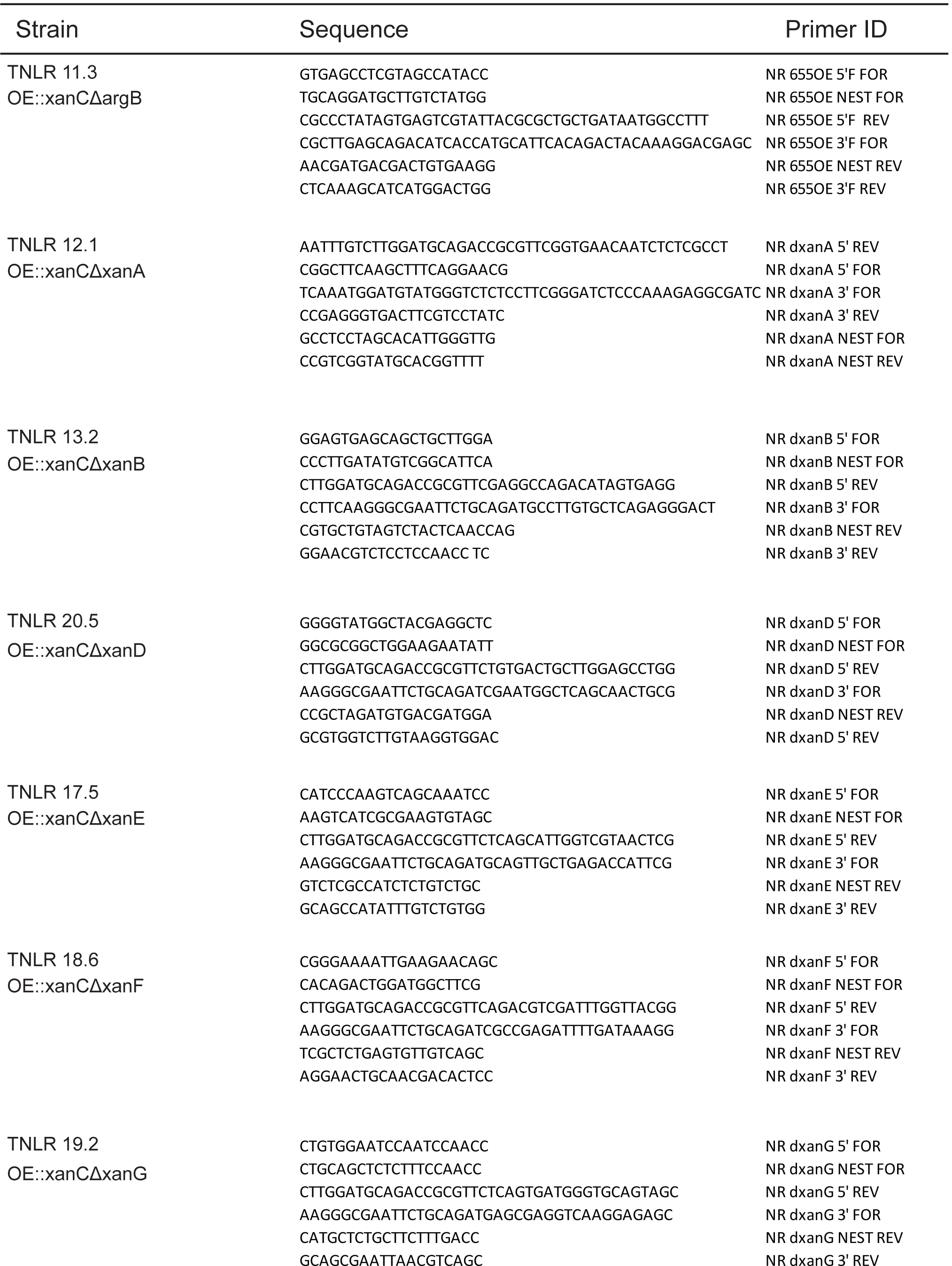
Primers used to generate fungal strains.

**Supplemental Table S3.**
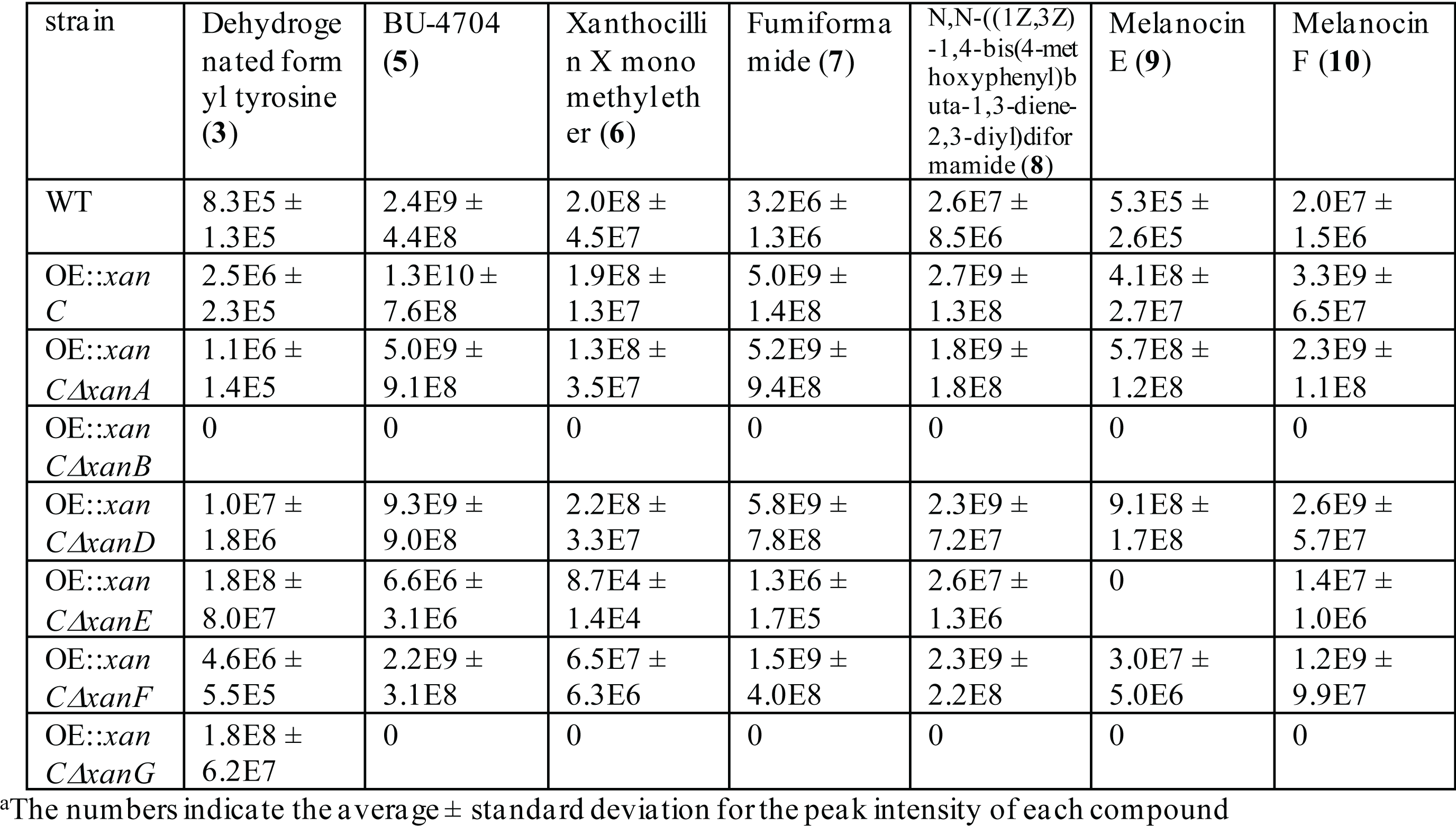
Production of xanthocillin derivatives comparing the OE::*xanC* mutants to deletion strains and WT.

**Supplemental Table S4.** UHPLC-HRMS data for dehydrogenated formyl tyrosine.

| Compound | HRESI(+/-) observed (*m/z*) | Ion | Calculated ion formula | Calculated *m/z* | Retention time (min) |
| --- | --- | --- | --- | --- | --- |
| Dehydrogenated formyl tyrosine (3) | 230.04240 | [M+Na]<sup>+</sup> | C<sub>10</sub>H<sub>9</sub>O<sub>4</sub>NNa<sup>+</sup> | 230.04238 | 3.7 |

**Supplemental Figure S3.**
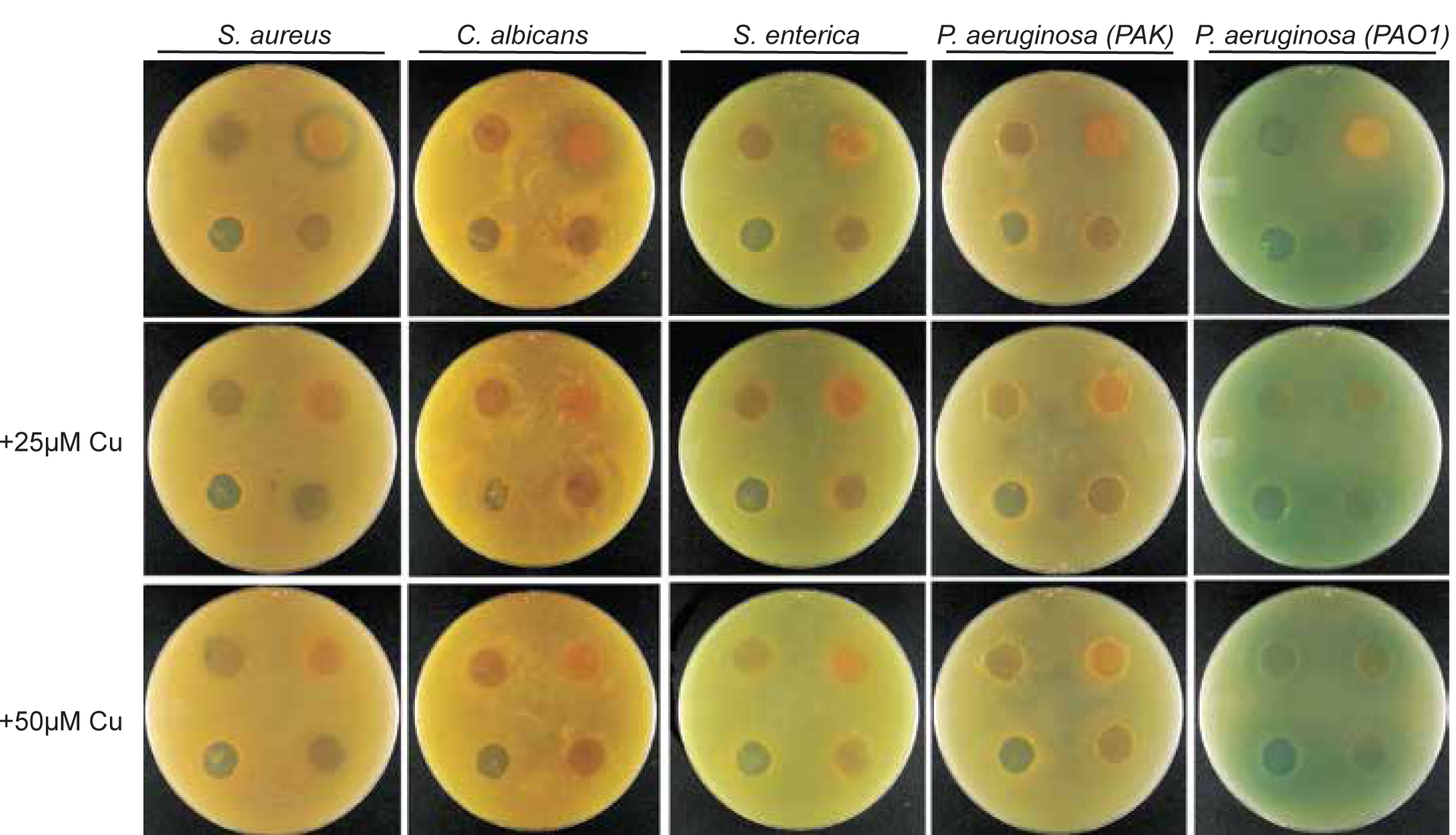
*xan* mutant antimicrobial assay. Agar plugs of actively growing *xanC* mutrants were transferred to a plate of lysogeny agar containing the indicated microbe (Top of figure) and supplemented with copper (Left of figure). Per-plate: Top left-WT, top right - OE::*xanC*, bottom left Δ*xanC*, bottom right-negative control (GMM).

**Supplemental Figure S4.**
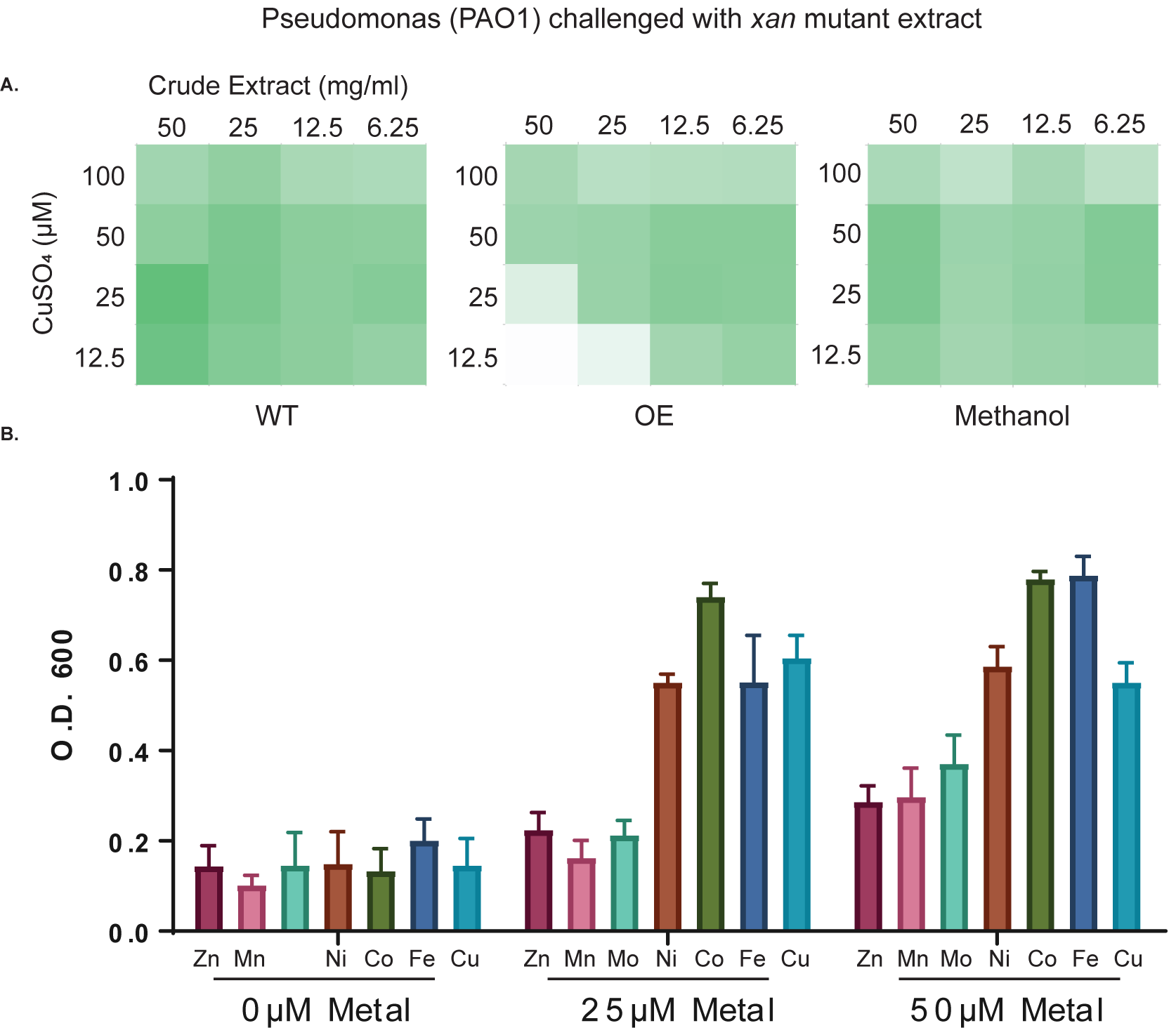
*Pseudomonas aeruginosa* PAO1challenged with *xan* mutant extracts. A.) Heat map indicating the relative growth of *P. aeruginosa* when challenged with crude extract (x-axis) and supplemented with copper (y-axis). Wild type, OE::*xanC* extract, or negative control is indicated at the bottom of the map. B.) Response of *P. aeruginosa* challenged with OE::*xanC* extract and supplemented with metals. Shown is the mean of three replicates with the SEM indicated by the error bars.

**Supplemental Figure S5.**
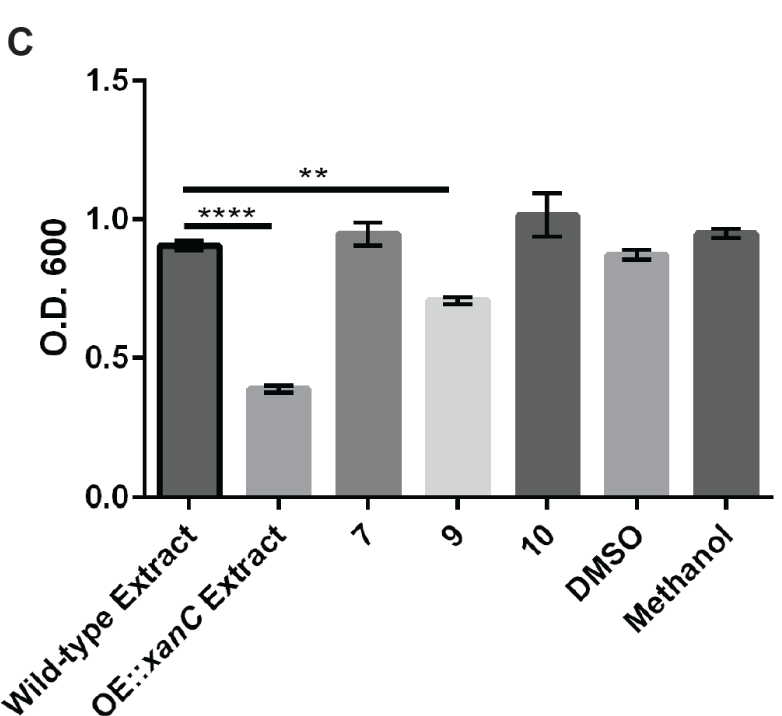
*Pseudomonas aeruginosa* PAO1 challenged with xanthocillin derivatives (256 μg/ml). Numbers are associated with the compounds in Figure 2. Shown is the mean of three technical replicates with the SEM indicated by the error bars. Statistical analysis was performed using an ANOVA. **P<0.01, ****P<0.0001.

**Supplemental Figure S6.**
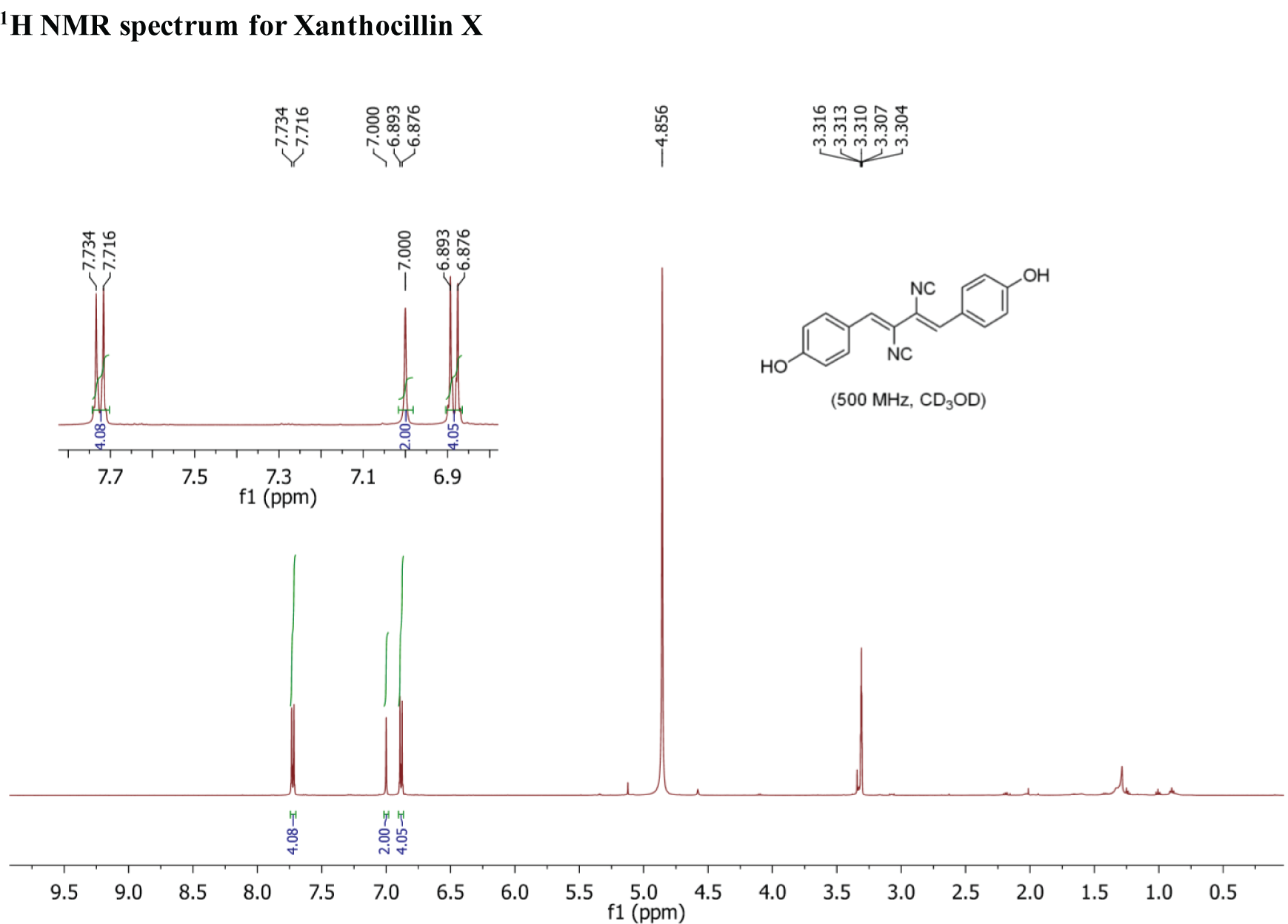
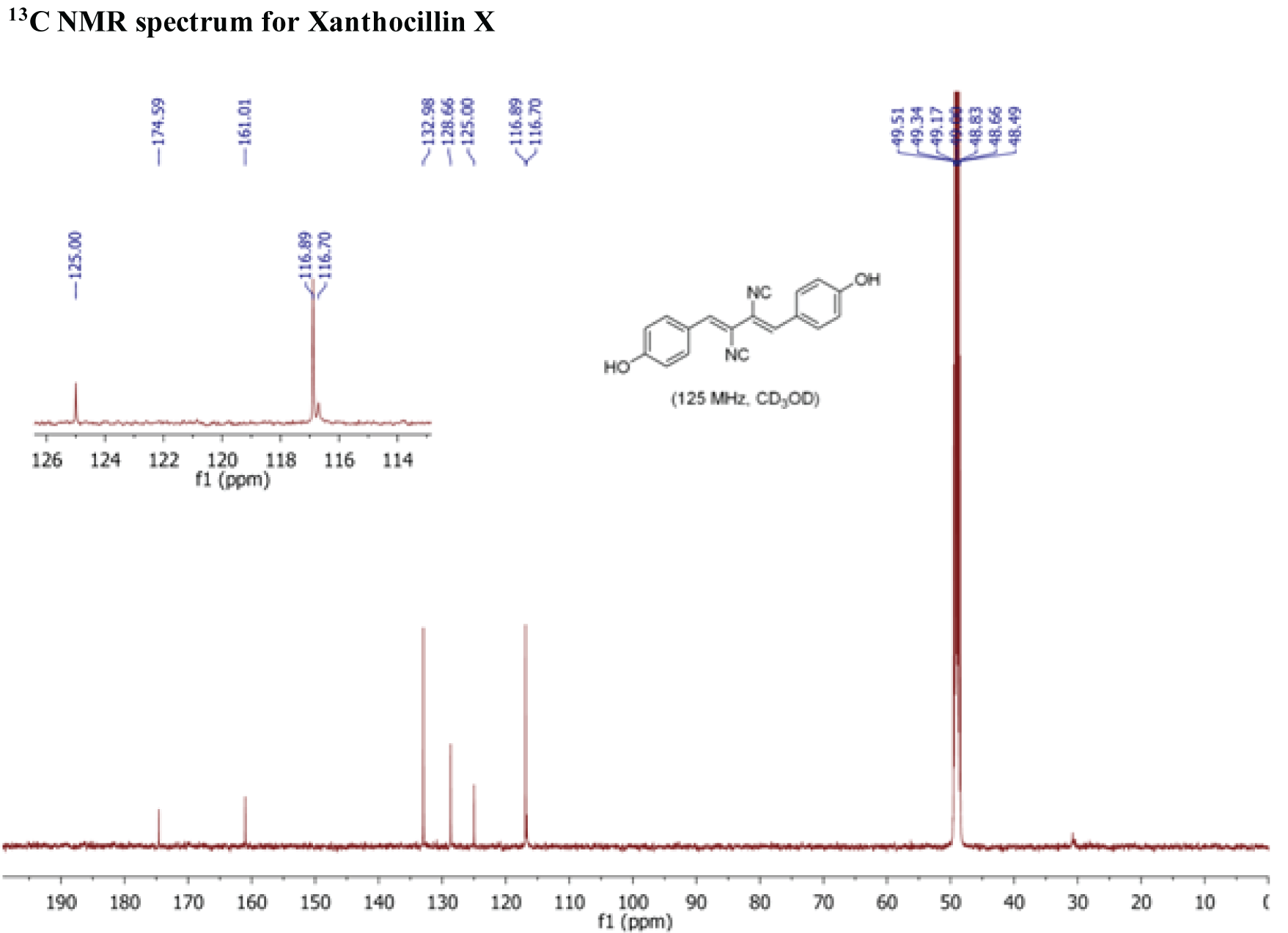
^1^H and ^13^C NMR spectra of synthetic xanthocillin.

